# Aerosolized ⍺_1_ adrenoreceptor antagonism does not affect experimentally induced lung fibrosis in animal models

**DOI:** 10.1101/2023.12.12.571378

**Authors:** Alexander Ghincea, Carrighan Perry, Angela Liu, John McGovern, Sheeline Yu, Xue-Yan Peng, Genta Ishikawa, Thomas Barnthaler, Erica L. Herzog, Huanxing Sun

**Author notes:** Corresponding author: (HXS).

## Abstract

Pulmonary Fibrosis is a progressive and incurable condition that complicates many disease states. Adrenergic hyperinnervation and accumulation of fibroblasts expressing ⍺_1_- adrenoreceptors have been implicated in this process. Previous studies have demonstrated that systemic treatment with an ⍺_1_-adrenoreceptors antagonist attenuates fibrotic endpoints in lung fibrosis models. In an attempt to develop a lung targeted therapy, we determined whether ⍺_1_- adrenoreceptors antagonism delivered via inhaled administration of terazosin exerts antifibrotic benefits in experimentally induced lung fibrosis. C57/BL6 mice treated with bleomycin, or a doxycycline inducible line of transgenic mice with lung specific overexpression of the bioactive form of the human TGFβ1 (*TGFβ1-Tg*^+^ model), received nebulized terazosin at varying doses on a therapeutic schedule following the induction of fibrosis and were sacrificed at 21 days. Airway inflammation, fibrotic endpoints, and lung function were evaluated. ⍺_1_-adrenoreceptors antagonism delivered via this method did not impact airway inflammation as indicated by bronchoalveolar lavage cell counts, and there was no significant difference observed in soluble collagen content. There was similarly no significant difference in respiratory mechanics with terazosin administration. These data show that inhaled delivery of the ⍺_1_-adrenoreceptors antagonist terazosin by this method is ineffective at treating fibrosis in these models and suggest that alternative dosing schedules or delivery methods may be more fruitful avenues of investigation. Further exploration of these findings may provide new therapeutic options and illuminate mechanisms through which adrenergic innervation and ⍺_1_-adrenoreceptors mediate fibrosis in the adult mammalian lung.

## Introduction

Pulmonary fibrosis (PF) is an incurable condition that complicates numerous interstitial lung diseases and is characterized by progressive pulmonary fibrosis and lung function decline [1, 2]. The incidence and demographic distribution of PF is heterogenous depending on underlying diagnosis but tends to occur in older adults and is often associated with a poor prognosis [3]. Idiopathic pulmonary fibrosis (IPF), one of the better studied forms of PF, is defined by a pattern of usual interstitial pneumonia on imaging and histopathology in the absence of a known cause and has a median survival of 2-4 years [4, 5]. Though the underlying pathologic mechanisms vary, pulmonary fibrosis is thought to be the result of dysregulated lung healing that results in the accumulation of activated fibroblasts and excessive extracellular matrix [6–8]. Despite numerous recent advances, treatments remain limited and lung transplantation is the only cure [4, 9]. Thus, considering the significant morbidity and mortality caused by these diseases, there is an urgent need for new targeted interventions to reduce the burden of fibrosis.

Noradrenergic mechanisms are increasingly associated with numerous pathologic processes including acute inflammation [10, 11], innate immune activation [12, 13], cancer [14], obesity and metabolism [15], bone marrow aging [16], and pathologic lung remodeling [17–19]. Adrenergic hyperinnervation and the accumulation of ⍺_1_-adrenoreceptor (⍺_1_-AR) expressing cells are associated with fibrosis in animal models, where interruption of adrenergic signaling via chemical denervation and ⍺_1_-AR antagonism mitigates fibrotic endpoints [17, 19]. It has further been shown that in some settings the blood [20] and lungs [19] of IPF patients are enriched for noradrenaline, and that IPF patients prescribed ⍺_1_-AR antagonists for non-pulmonary indications experience improved survival [19]. However, despite the increasing association of noradrenaline and fibrotic lung remodeling, the mechanisms of this effect and its therapeutic potential remain unexplored [21, 22].

Pulmonary expression of ⍺_1_-ARs is the best characterized in smooth muscle cells of the airways and vasculature, where they are reported to mediate noradrenaline-induced contractile responses [23, 24]. Interestingly, this receptor class is also expressed in parenchymal endothelium and fibroblasts [25], which may, in part, account for the capillary leak-associated fibroblast activation and extracellular matrix expansion caused by intravenous infusion of the ⍺_1_- AR agonist phenylephrine [26, 27] and the therapeutic benefit of systemically administered ⍺_1_-AR antagonism delivered via the oral or intraperitoneal routes [17–19]. However, ⍺_1_-ARs are also expressed by populations of lung epithelia and airway inflammatory cells which may contribute to noradrenaline’s well described role in remodeling responses of apoptosis [28], inflammatory cell trafficking [29], and cytokine production [30]. This expression pattern suggests that interventions leveraging inhaled administration might exert therapeutic benefit while minimizing off-target effects. Elucidation of this biology has the potential to provide insights into the cell(s) and mechanism(s) through which noradrenaline exerts its fibrogenic effects while exploring new treatment modalities [31–33] for parenchymal disease.

This study sought to address this knowledge gap by assessing whether inhaled administration of the non-selective ⍺_1_-AR antagonist terazosin reduces lung inflammation, pulmonary function, biochemical collagen accumulation, and lung histology in two well accepted animal models of experimentally induced fibrosis.

## Materials and Methods

### Study Design

The objective of our study was to test the prespecified hypothesis that nebulized delivery of an α1-AR antagonist, specifically terazosin, would improve experimentally induced pulmonary fibrosis in two commonly used mouse models. Primary endpoints were prospectively determined.

### Animals

All animal experiments were approved by the Yale School of Medicine IACUC in accordance with federal regulations (protocol #20292). Twenty-one days following the experimental induction of fibrosis, animals were anesthetized for pulmonary function testing, then humanely sacrificed to isolate bronchoalveolar lavage (BAL), and *en bloc* resection of lungs for soluble collagen quantification and histopathology scoring as we have previously described [34]. C57 Black 6 (C57BL/6) wild-type mice were used for bleomycin experiments. TGFβ1 lung specific over-expressing (*CC10-tTS-rtTA-TGFβ1,* from here on called “*TGFβ1-Tg*^+^”) mice have been described previously [35] and develop fibrosis in response to doxycycline inducible expression of the bioactive form of the human *TGFβ1* gene under the lung specific CC10 promoter. All experimental groups were matched for age and sex.

### Bleomycin model

Sex-matched C57BL/6 aged 9 to 11 weeks mice were exposed to a single dose of 1.5 U/kg pharmacologic grade bleomycin (Northstar Rx LLC, NDC 16714-886-0) or sterile saline by orotracheal aspiration [36]. Mice were anesthetized with isoflurane and suspended by their incisors on a standing rack. With the tongue held in gentle retraction, 50 µL bleomycin or saline was pipetted into the oropharynx and aspirated. Animals were sacrificed after 21 days and evaluated for the endpoints described above.

### TGFβ1-Tg^+^ model

Sex-matched 9 to 11-week-old wild-type *TGFβ1-Tg^+^* mice received 0.5 mg/mL doxycycline hyclate (Thermo Scientific Cat No. J60579.22) in their drinking water for 21 days as previously described [35].

### Nebulization Experiments

Starting at day 5 after bleomycin administration or doxycycline initiation, mice received daily administration of nebulized terazosin (Millipore Sigma, Product No. 1643452) dissolved in sterile water at concentrations of 0.001 mg/mL, 0.002 mg/mL, 0.01 mg/mL, 0.02 mg/mL, 0.1 mg/mL, and 0.2 mg/mL which were derived to approximate a total inhaled dose of 0.001 mg/kg,

0.002 mg/kg, 0.01 mg/kg, 0.02 mg/kg, 0.1 mg/kg, and 0.2 mg/kg per day (Fig S1) [37].

Concentrations that showed potential benefit were selected for confirmatory studies. Groups of up to 5 mice from a single cage were placed in a clean plastic box with an attachment for nebulizer tubing in a manner similar to that used by Schroeder et al [38] for drug administration. Omron CompAir NE-C28 compressor nebulizers were used to aerosolize and deliver 7 mL of terazosin or sterile water control on a daily therapeutic schedule over 30 min after which mice were returned to their original cages. The same terazosin dose was repeated for every treatment within each group. A schematic of the nebulizer apparatus is shown in Fig 1A.

**Fig 1:**
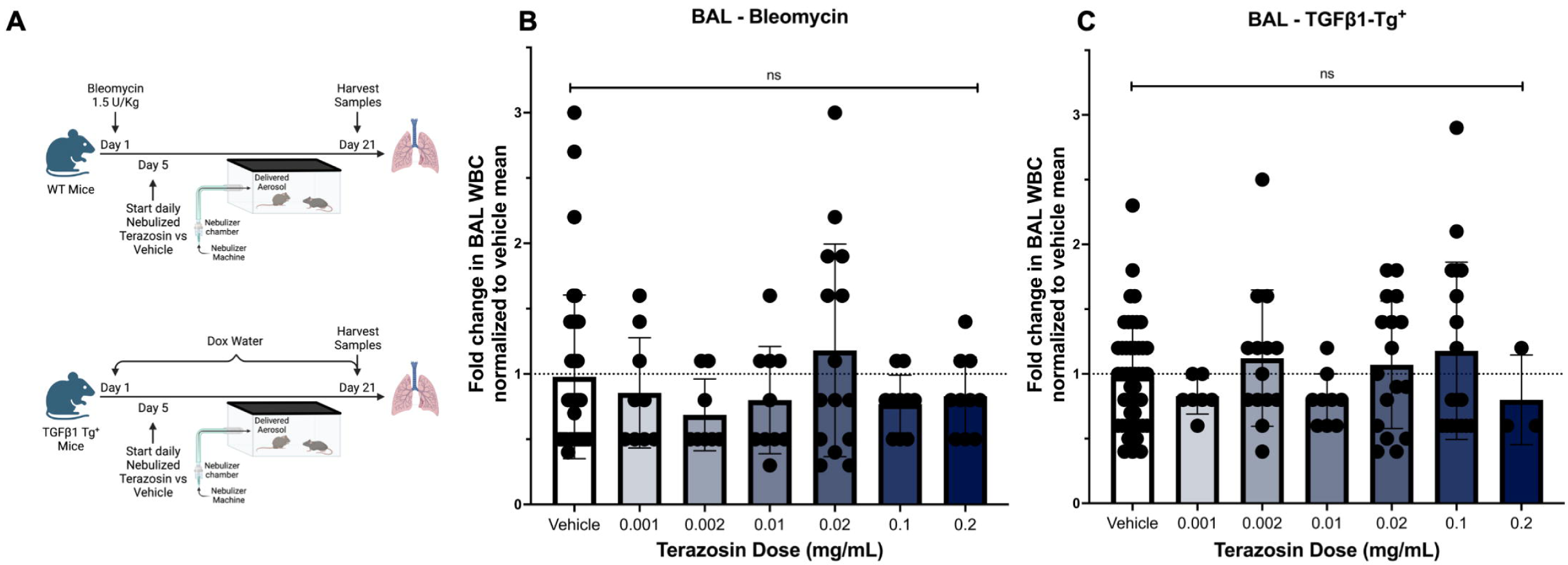
Inhalational administration of the nonselective ⍺_1_-AR antagonist terazosin does not impact lung inflammation. *A*: Experimental design. Inhaled terazosin was started five days after bleomycin administration or doxycycline induced TGFβ1 transgene activation. Up to 5 mice from a single cage were placed into a clean, lidded plastic box with an attachment for nebulizer tubing. 7 mL of sterile water or terazosin was loaded into the nebulizer module for each treatment. Tubing connections were checked to be airtight prior to every treatment. Omron NE-C28 compressor nebulizers were used to aerosolize and deliver the treatment over 30 minutes. After treatment mice were returned to their original cages, and the plastic box and nebulizer equipment cleaned thoroughly. ***B***: In the bleomycin model, relative to vehicle, inhaled terazosin had no significant impact on lung inflammation as measured by BAL cell counts. ***C***: In the TGFβ1-Tg^+^ model, relative to vehicle, inhaled terazosin had no significant impact on lung inflammation as measured by BAL cell counts. Schematic diagrams created using BioRender.com

### BAL cell count

At the time of sacrifice, two aliquots of 0.8 mL of PBS were slowly instilled into lung and the lavage fluid gently aspirated. The combined BAL sample was assessed for white blood cell counts using a Beckman Coulter Ac.T Diff instrument.

### Pulmonary function testing

Pulmonary mechanics and airway hyperresponsiveness were assessed in a subset of terazosin-exposed and control mice from both bleomycin and *TGFβ-Tg^+^* arms of our study using the Sireq FlexiVent Fx computer-controlled piston ventilator running Flexiware version 7.6 [39]. Animals were anesthetized with an intraperitoneally administered mixture of xylazine and ketamine, xylazine (Akorn, NDC 59399-110-20) at a dose of 10 mg/kg and ketamine (Covetrus, NDC 11695-0703-1) at a dose of 100 mg/kg, followed by urethane at a dose between 0.009 to 0.03 g per mouse, administered intraperitoneally. Once an appropriate level of sedation was established, a midline neck incision was made, and the soft tissue dissected to expose the trachea. A rigid 20-guage blunt tipped catheter for female mice or 18-gauge blunt tipped catheter for male mice was used to cannulate the trachea. Mice were then paralyzed with pancuronium bromide (Sigma, Cat# P1918-10mg) at a dose of 10 µg per mouse and attached to the FlexiVent for pulmonary function measurements. Methacholine (SigmaAldrich A2251-25G) was used for airway hyperresponsiveness measurements. After completion of lung function testing, mice were humanely euthanized and harvested for tissue as described above.

### Collagen quantification

Right lung soluble collagen content was assessed using the Sircol Soluble Collagen Assay (Biocolor Ltd., CLS 1111) according to the manufacturer’s instructions.

### Histologic analysis

Formalin fixed and paraffin embedded (FFPE) tissue sections obtained from whole left lungs harvested from experimental animals were stained with Masson’s trichrome stain to visualize collagen deposition. Modified Ashcroft Scores, a semiquantitative scoring system validated for experimentally induced fibrosis in rodents, were also determined [40].

### Statistics

GraphPad Prism version 9.4.1 was used for statistical analysis and data graphing. Unpaired 2-tailed Mann-Whitney and Kruskal-Wallis tests were used for nonparametric comparisons, and 2-tailed Students T-test and Analysis of Variance used for parametric comparisons. The slopes of expiratory flow-volume loops and airway resistance curves were compared using simple linear regression.

## Results

### Inhalational administration of an ⍺_1_-AR antagonist does not impact lung inflammation

⍺_1_-AR antagonists administered via oral or intraperitoneal routes demonstrate antifibrotic benefit in models of cross organ fibrosis including the lung, liver, heart, and kidney [41–44]. To determine whether similar benefit is achieved by local delivery to the lung, the following experiments were performed. Because the benefits of ⍺_1_-AR antagonism include attenuation of inflammation [45, 46] and TGFβ1-dependent tissue responses [47], two experimental models were used: single dose inhaled bleomycin, and lung specific, doxycycline inducible overexpression of the bioactive form of the human *TGFβ1* gene. In both models, mice received nebulized terazosin according to a therapeutic schedule (Fig 1A). Doses were determined based on previous literature indicating a 1:1 conversion between systemic and inhaled delivery of the related ⍺_1_-AR antagonist, Prazosin [48]. Because ⍺_1_-AR antagonism mitigates lung inflammation in a number of contexts, we began by determining whether this treatment might reduce the airway inflammation associated with pulmonary fibrosis in the hyperinflammatory bleomycin mode. The assessment of BAL cell counts in the bleomycin model revealed no significant change following administration of nebulized terazosin at any dose (Fig 1B). Similar findings were seen in the *TGFβ1-Tg*^+^ model, where BAL cell counts remained essentially unaltered by terazosin (Fig 1C). These data indicate that at the range of doses tested, aerosolized delivery of an ⍺_1_-AR antagonist does not impact the magnitude of lung inflammation.

### Inhalational administration of an ⍺_1_-AR antagonist does not impact lung compliance or airway resistance

Physiologic changes caused by pulmonary fibrosis include the development of restrictive lung physiology and our previous work has shown that patients prescribed ⍺_1_-AR antagonists for prostate-related indications exhibited preserved lung compliance as reflected by the forced vital capacity (FVC) [19] and could improve lung compliance without a concomitant increase in airway resistance that could be theoretically induced by ⍺_1_-AR signaling [48]. To test this hypothesis, bleomycin-challenged mice that had or had not received inhaled terazosin underwent pulmonary function testing using the FlexiVent system. In the bleomycin model, mean baseline static lung compliance of 0.046 mL/cmH2O (Fig 2A), pressure-volume loops (Fig 2B), and mean baseline maximal airway resistance of 2.96 cmH2O.s/mL assessed following methacholine challenge (Fig 2C) were not significantly changed by terazosin administration. Similar findings were observed in the *TGFβ1-Tg^+^* mice, where baseline static compliance of 0.044 mL/cmH2O and airway resistance of 5.66 cmH2O.s/mL did not display changes in response to nebulized terazosin (Fig 2D-F). These data show that aerosolized delivery of the ⍺_1_-AR antagonist terazosin is ineffective in altering lung physiology in two models of fibrotic ILD.

**Fig 2:**
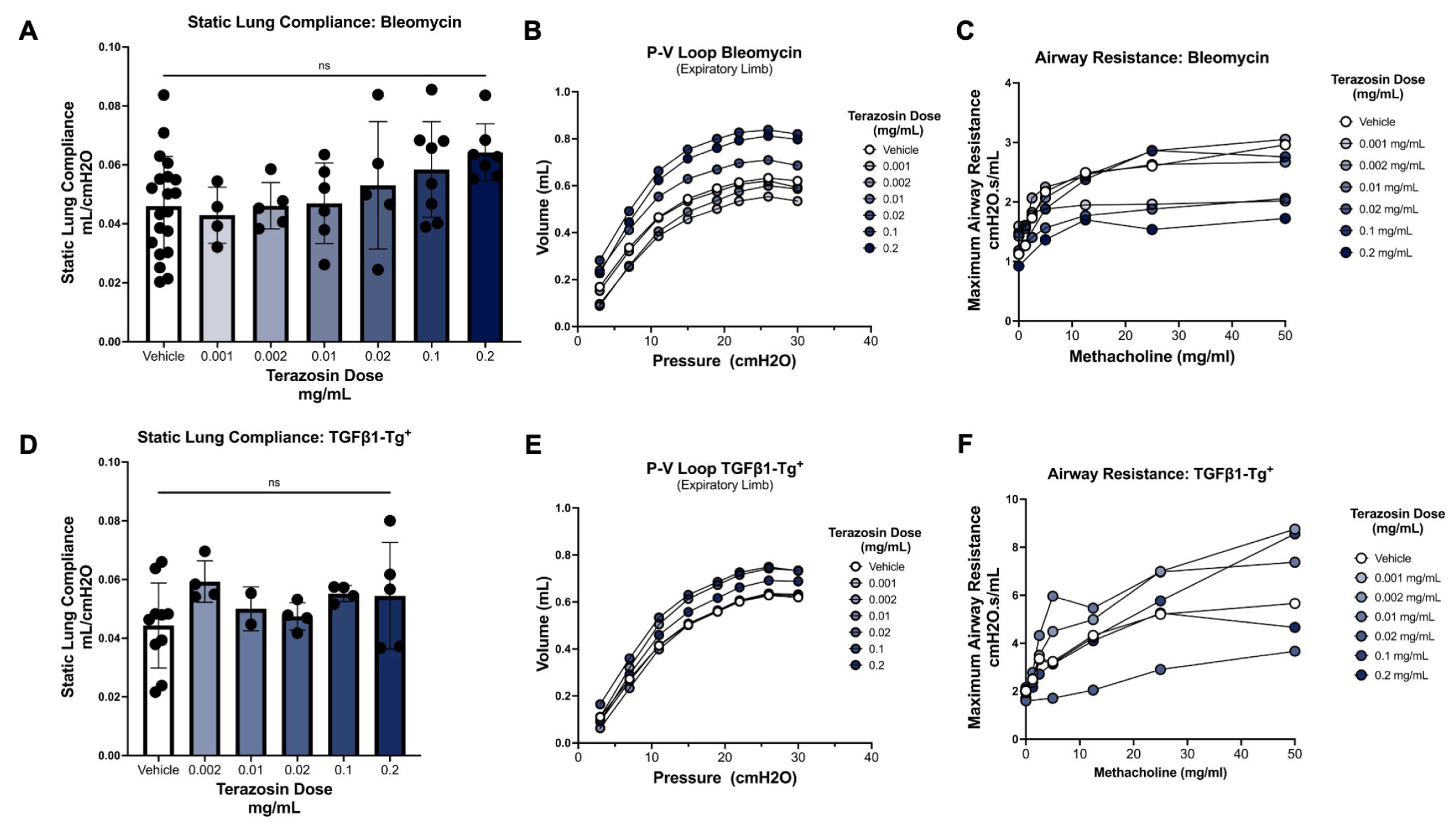
Inhalational administration of an ⍺_1_-AR antagonist does not impact lung compliance or airway resistance. *A*: Bleomycin-challenged mice that had or had not received inhaled terazosin underwent pulmonary function testing using the FlexiVent system. Here, a mean baseline static lung compliance of 0.046 mL/cmH2O was observed in untreated mice and was not significantly altered by terazosin administration. ***B***: Terazosin did not significantly change the pressure-volume relationship for lungs tested. ***C***: Maximal airway resistance assessed following methacholine challenge did not significantly change following terazosin administration from a mean baseline of 2.96 cmH2O.s/mL in untreated mice. ***D-F***: Similar findings were observed in the TGFβ1-Tg^+^ mice, where baseline static compliance of 0.044 mL/cmH2O and airway resistance of 5.66 cmH2O.s/mL did not display changes in response to nebulized terazosin. These data show that aerosolized delivery of the ⍺_1_-AR antagonist terazosin is ineffective in altering lung physiology in two models of fibrotic ILD.

### Inhalational administration of an ⍺_1_-AR antagonist does not impact lung collagen accumulation

Systemic administration of ⍺_1_-AR antagonists has been shown to suppress fibroblast activation and its associated accumulation of collagen in numerous models of organ fibrosis. Our own work and that of others shows that this effect extends to the lungs [17, 19]. Reasoning that local delivery of the ⍺_1_-AR antagonist terazosin might show similar efficacy, the lungs of bleomycin-challenged mice that did and did not receive terazosin underwent biochemical quantification of collagen using the well described Sircol assay. Using this approach, fold change from a baseline collagen content per whole right lung was essentially unchanged across all terazosin doses tested (Fig 3A). In the *TGFβ1-Tg^+^* model, promising reductions in the fold change from baseline collagen content per whole right lung were observed at several tested terazosin doses (Fig 3B). Based on these results we selected the terazosin doses of 0.002 and 0.02 mg/mL for confirmatory studies but were unable to replicate our initial results (Fig 3C-D). Given these conflicting findings, we sought to confirm our results using histological measures such as trichrome staining and semi-quantitative modified Ashcroft scores, which complement biochemical measures by assessing collagen accumulation in parenchymal regions. The results of these studies were consistent with our confirmatory biochemical measurements, as in the *TGFβ1-Tg^+^* model trichrome staining and baseline modified Ashcroft scores was similar at the doses of terazosin assessed (Fig 3E-F). When viewed in combination, our data show that localized delivery of aerosolized terazosin exerts no observable benefit in these models and is insufficient to mitigate lung fibrosis.

**Fig 3:**
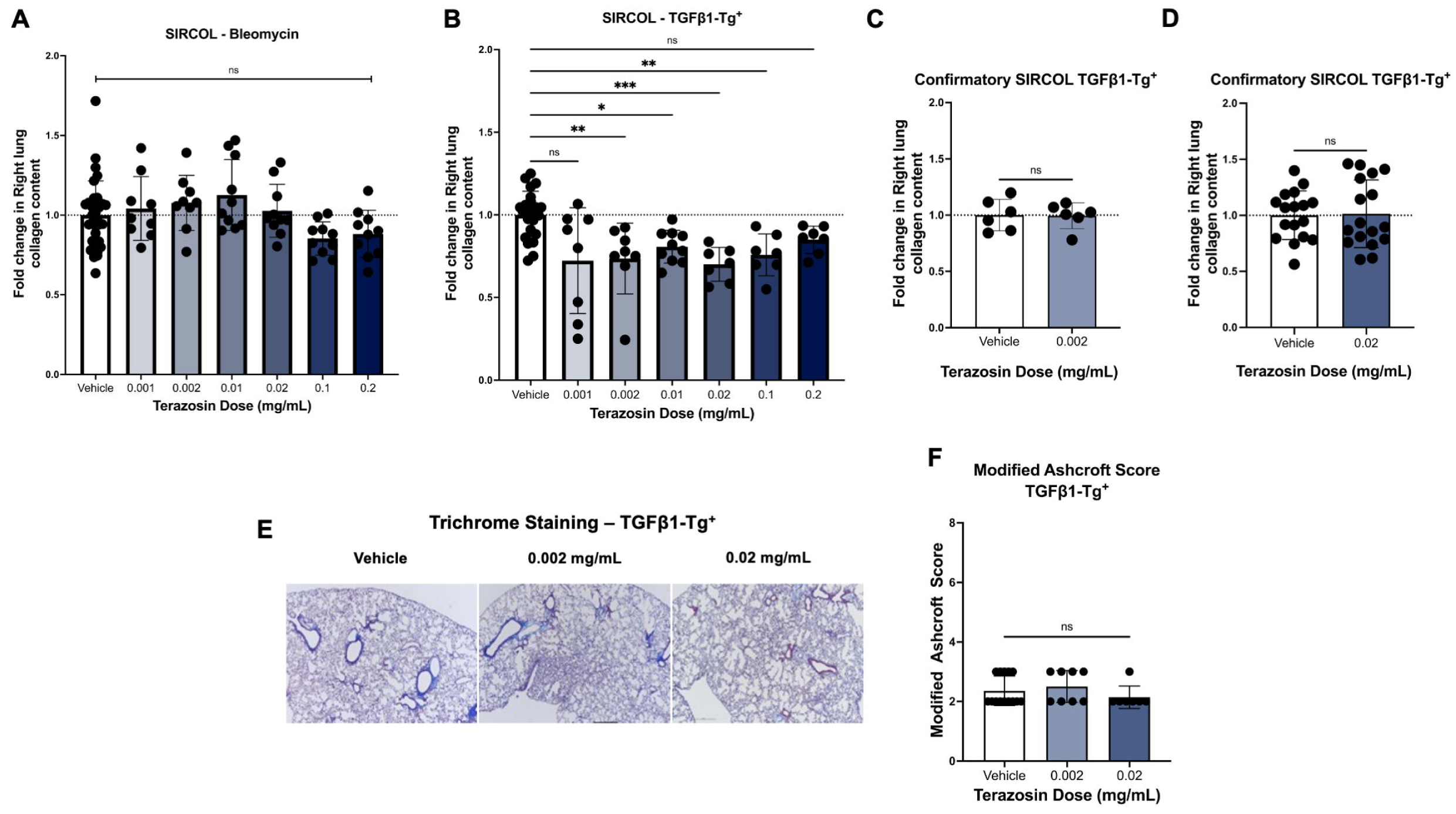
Inhalational administration of an ⍺_1_-AR antagonist does not impact lung collagen accumulation. ***A***: Fold change from baseline collagen content per whole right lung was essentially unchanged across all doses tested in bleomycin challenged mice. ***B***: In the TGFβ1- Tg^+^ model, initial studies revealed a significant difference in fold change from baseline right lung collagen content at 0.002 mg/mL, 0.01 mg/mL, 0.02 mg/mL, and 0.1 mg/mL Terazosin. ***C-D***: Due to these initially promising results confirmatory studies were performed for 0.002 mg/mL and 0.02 mg/mL. We were unable to replicate the initially promising differences seen in TGFβ-Tg^+^ animals at Terazosin doses of 0.002 mg/mL and 0.02 mg/mL. ***E-F***: Trichrome staining and semi-quantitative Modified Ashcroft Scores done in our confirmatory studies did not significantly change after administration of terazosin. These data show that in contrast to the previously reported antifibrotic benefit of systemic ⍺_1_-AR, nebulized delivery of this agent is insufficient to mitigate collagen deposition in lung tissues.

## Discussion

This study aimed to determine whether local delivery of the ⍺_1_-AR antagonist terazosin improves markers of pulmonary fibrosis in two different animal models. Our data indicate that the administration of terazosin in nebulized form did not impact lung inflammation, airway compliance or resistance, collagen accumulation, or histologic measures of lung remodeling in the bleomycin and *TGFβ1-Tg^+^* models of murine pulmonary fibrosis. These results are in contrast to prior work in which intraperitoneal administration of terazosin modulates fibrotic changes in bleomycin challenged mice [17] and numerous studies demonstrating that ⍺_1_-AR blockade improves fibrotic endpoints in cross-organ models of tissue fibrosis including the lung, liver, heart, and kidney [41–44]. Notably, because the present study assessed localized tissue delivery as opposed to systemic ⍺_1_-AR antagonism, our data indicate that alternate approaches will be required to support the repositioning of this drug for the treatment of lung fibrosis.

The ⍺_1_-ARs are a conserved family of G-protein coupled receptors found in numerous tissues including the lung and have been identified both intracellularly and on the extracellular membrane of smooth muscle cells in the airway and pulmonary vasculature, immune cells, and fibroblasts [23, 24, 49]. Additionally ⍺_1_-AR agonism has been linked to increased expression of inflammatory markers such as IL-1⍺, IL-1β, IL-6, and TNF-⍺ both systemically and in the lungs, and prior work has associated this to increased levels of TGFβ [27]. In contrast, systemically delivered ⍺_1_-AR antagonism has been shown to attenuate lung inflammation [45, 46] and TGFβ1- dependent tissue responses [47], is associated with improved fibrotic endpoints [41–44] and survival in IPF patients [19]. One interpretation of our conflicting results is that administration of ⍺_1_-AR antagonist by nebulized aerosol might only interact with cells locally in the airway and alveolus and may not achieve sufficient uptake to the fibrotic interstitium to exert its effects on the fibroblasts, myofibroblasts, and pericytes that are the primary effectors of fibrosis (Fig 4).

**Fig 4:**
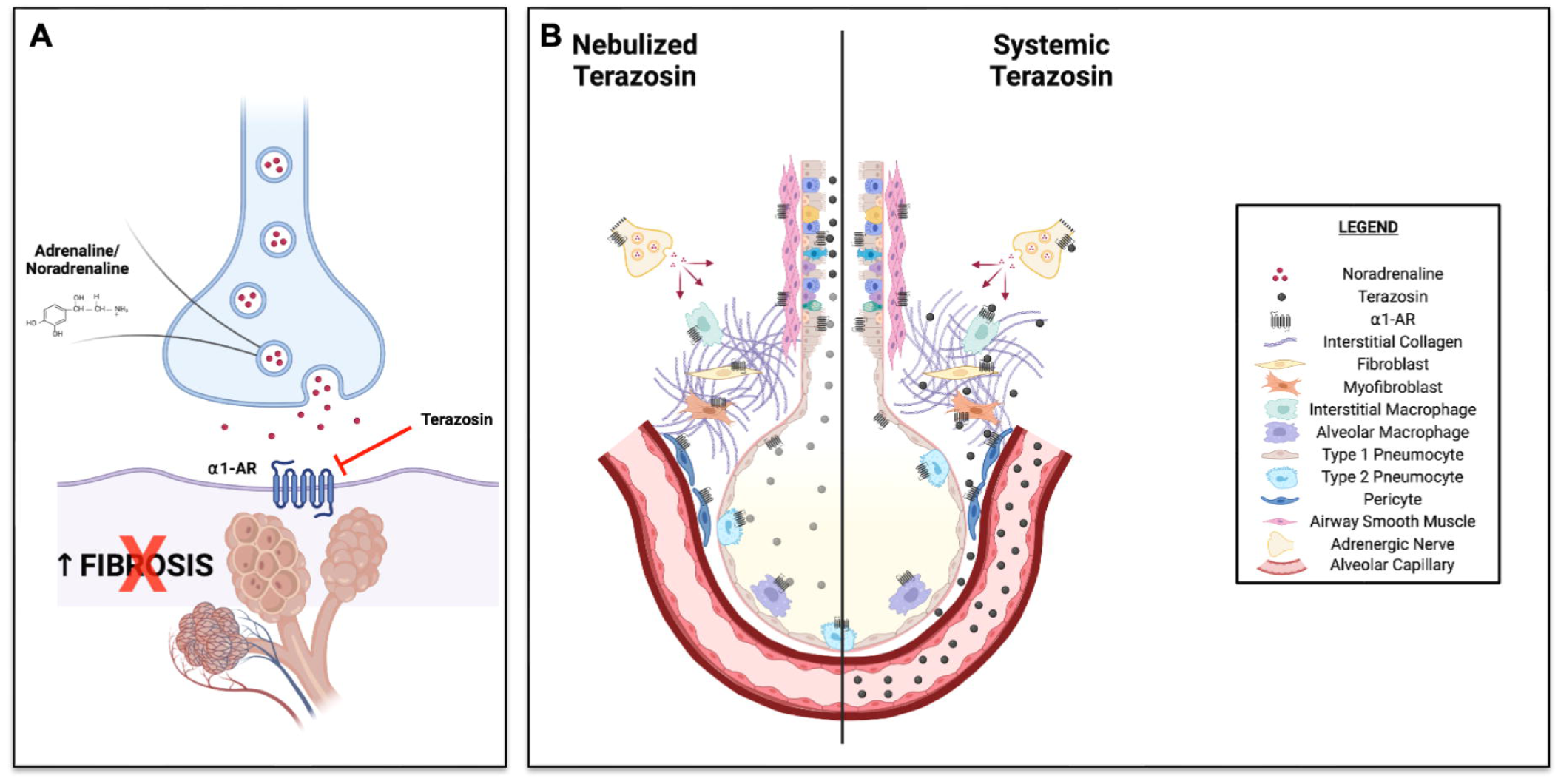
Proposed schematics. ***A***. Schematic of the proposed basic mechanism of terazosin. By non-selectively antagonizing ⍺_1_-ARs, Terazosin is proposed to block the pro-fibrotic effects of noradrenaline in the lung. ***B***. Adrenergic nerve density is increased in the fibrotic lung and increased noradrenaline is believed to promote a fibrotic phenotype. Numerous cell types including fibroblasts, myofibroblasts, macrophages, airway epithelial cells, and sympathetic neurons express ⍺_1_-ARs, and these are up-regulated in fibrosis. There are several mechanisms which may explain the lack of anti-fibrotic effect observed in our study. When terazosin is nebulized and inhaled, particulate size, dosage, and the heterogeneous nature of fibrotic lung may limit the amount of drug that reaches the terminal airways and alveoli (indicated by more lightly shaded particulates in the schematic). Additionally, though remodeling responses secondary to noradrenaline have been described in the epithelium, the terazosin dose when administered in nebulized form may be insufficient to alter pro-fibrotic signaling. Alveolar macrophages, which are more likely to be exposed to terazosin, may not play as large a role in the development of fibrosis in our model. Finally, transport of terazosin across the basement membrane may also be limited so that insufficient quantities of the drug reach interstitial fibroblasts, myofibroblasts, and pericytes. In contrast, systemic administration of terazosin is more likely to achieve therapeutic levels for ⍺_1_-AR antagonism in fibrotic foci. Created with BioRender.com

Terazosin is a specific ⍺_1_-AR antagonist [50] that has been approved for use for decades in the treatment of benign prostatic hyperplasia and hypertension, and has a well-established safety profile [51, 52]. Its long half-life makes it well suited for once daily administration [53]. Though there are no prior studies assessing inhalational forms of terazosin, other ⍺_1_-AR antagonists have been studied in the context of asthma [54] including a nebulized form of another long acting ⍺_1_-AR antagonist Prazosin [48, 55, 56]. While these early studies had limited efficacy in the overall management of asthma, they did indicate that nebulized delivery of ⍺_1_-AR antagonists is a viable modality. However, the fibrotic lung is spatially heterogenous with areas of normal tissue interspersed with diseased lung that has mismatched ventilation and perfusion [57]. Numerous agents are in development or have been investigated for topical delivery in IPF and recent work has demonstrated that particulate dimension is critically important for adequate drug delivery to fibrotic sites [58]. Specifically, Usmani and colleagues demonstrated that a particulate size of 1.5 µm compared to 6 µm was best for uptake in the fibrotic lung [58]. The Omron CompAir NE-C28 compressor nebulizers used in the present study produce an aerosol particle size of 3 µm median mass aerodynamic diameter. While suboptimal based upon the aforementioned study, we believe that this particle size should reach the terminal airways. However, particulate size may yet represent a potential source for the negative results observed in our study. If too large, the particulates may become trapped in the oropharynx, trachea, or larger airways and fail to reach fibrotic regions.

Our study is limited in not having assessed the spatial delivery of drug within the mouse lung nor systemic uptake of terazosin into the bloodstream. Target engagement was also not evaluated. Furthermore, our apparatus delivered the aerosolized terazosin into a box containing several mice such that at least some amount of the aerosol was deposited onto the fur of treated animals as well as the interior surface of the container, limiting the quantity of terazosin inhaled into the lungs. Oral ingestion of drug accumulated on the fur or interior surface may further confound our results. Finally, the dose of terazosin administered, or dosing schedule utilized may have been insufficient to achieve adequate penetration or ⍺_1_-AR blockade. Recent work by Bai et al using a nanoparticle delivery system presents an alternative method for blocking ⍺_1_-ARs that can be investigated in future studies [59].

## Conclusion

Aerosolized delivery of the ⍺_1_-AR antagonist terazosin is ineffective at treating fibrosis in the bleomycin and *TGFβ1-Tg*^+^ models of murine pulmonary fibrosis. Given existing data showing benefit from systemic ⍺_1_-AR antagonism in fibrotic models, alternative dosing schedules or modes of delivery may be more fruitful avenues of research. Further exploration of these findings may provide new therapeutic options and illuminate the mechanisms through which adrenergic innervation and ⍺_1_-ARs mediate fibrosis in the adult mammalian lung.

## Acknowledgements

This work was supported by The Assistant Secretary of Defense for Health Affairs endorsed by the Department of Defense, in the amount of ($334,999.00), through the Peer Reviewed Medical Research Program under Award Number W81XWH-20-1-0157. Opinions, interpretations, conclusions, and recommendations contained herein are those of the author(s) and are not necessarily endorsed by the Department of Defense. The U.S. Army Medical Research Acquisition Activity, 820 Chandler Street, Fort Detrick MD 21702-5014 is the awarding and administering acquisition office. In conducting research using animals, the investigator(s) adhere(s) to the laws of the United States and regulations of the Department of Agriculture. In the conduct of research utilizing recombinant DNA, the investigator(s) adhered to NIH Guidelines for research involving recombinant DNA molecules. In the conduct of research involving hazardous organisms or toxins, the investigator(s) adhered to the CDC-NIH Guide for Biosafety in Microbiological and Biomedical Laboratories.

We are grateful for Dr. Qing Liu’s instruction of the operation of the FlexiVent System and the Omron CompAir NE-C28 compressor nebulizer.

## Supporting information

**Fig S1**: **Method used to derive Terazosin doses administered. *A***: The time (Tf) to fill the box in which mice were placed for nebulization experiments with nebulized drug is represented by the quotient of Box Volume divided by the Nebulizer Flow Rate. Our enclosures have a volume of 6682 mL, and the Omron CompAir NE-C28 nebulizer has a flow rate of 4000 mL/min. ***B***: The aggregate rate (R) at which nebulized drug enters the box is equal to the total Volume of Drug used divided by the Nebulization Time multiplied by the Drug Concentration loaded into the nebulizer. 7 mL of drug was used in our experiments, and based on our observations, the Omron CompAir NE-C28 nebulizer requires approximately 30 minutes to nebulize all 7 mL. ***C***: The drug weight (Dw) of nebulized terazosin within our enclosure is equal to the product of Tf and R. ***D***: When divided by the box volume, this results in the nebulized drug’s concentration (Dc) within the box. Minute Ventilation (^V̇^ E) is equal to the product of Respiratory Rate and Tidal Volume. The respiratory rate for adult mice is 187 breaths per minute, and the tidal volume 0.061 mL and 0.094 mL for 20g and 25g mice respectively (38). ***E***: Thus, we derive the total dose of drug in milligrams inhaled (Dt) by a single mouse during each 30-minute nebulization treatment to be equal to the product of Dc, ^V̇^ E, and nebulization time.

## Notes

### Competing Interest Statement

The authors have declared no competing interest.

### Summary of Updates

In the Acknowledgement, we added below information. This work was supported by The Assistant Secretary of Defense for Health Affairs endorsed by the Department of Defense, in the amount of ($334,999.00), through the Peer Reviewed Medical Research Program under Award Number W81XWH-20-1-0157. Opinions, interpretations, conclusions, and recommendations contained herein are those of the author(s) and are not necessarily endorsed by the Department of Defense. The U.S. Army Medical Research Acquisition Activity, 820 Chandler Street, Fort Detrick MD 21702-5014 is the awarding and administering acquisition office. In conducting research using animals, the investigator(s) adhere(s) to the laws of the United States and regulations of the Department of Agriculture. In the conduct of research utilizing recombinant DNA, the investigator(s) adhered to NIH Guidelines for research involving recombinant DNA molecules. In the conduct of research involving hazardous organisms or toxins, the investigator(s) adhered to the CDC-NIH Guide for Biosafety in Microbiological and Biomedical Laboratories.

