## Supplementary figures and images for "Aerosolized ⍺_1_ adrenoreceptor antagonism does not affect experimentally induced lung fibrosis in animal models"

### Supplemental Figure 1

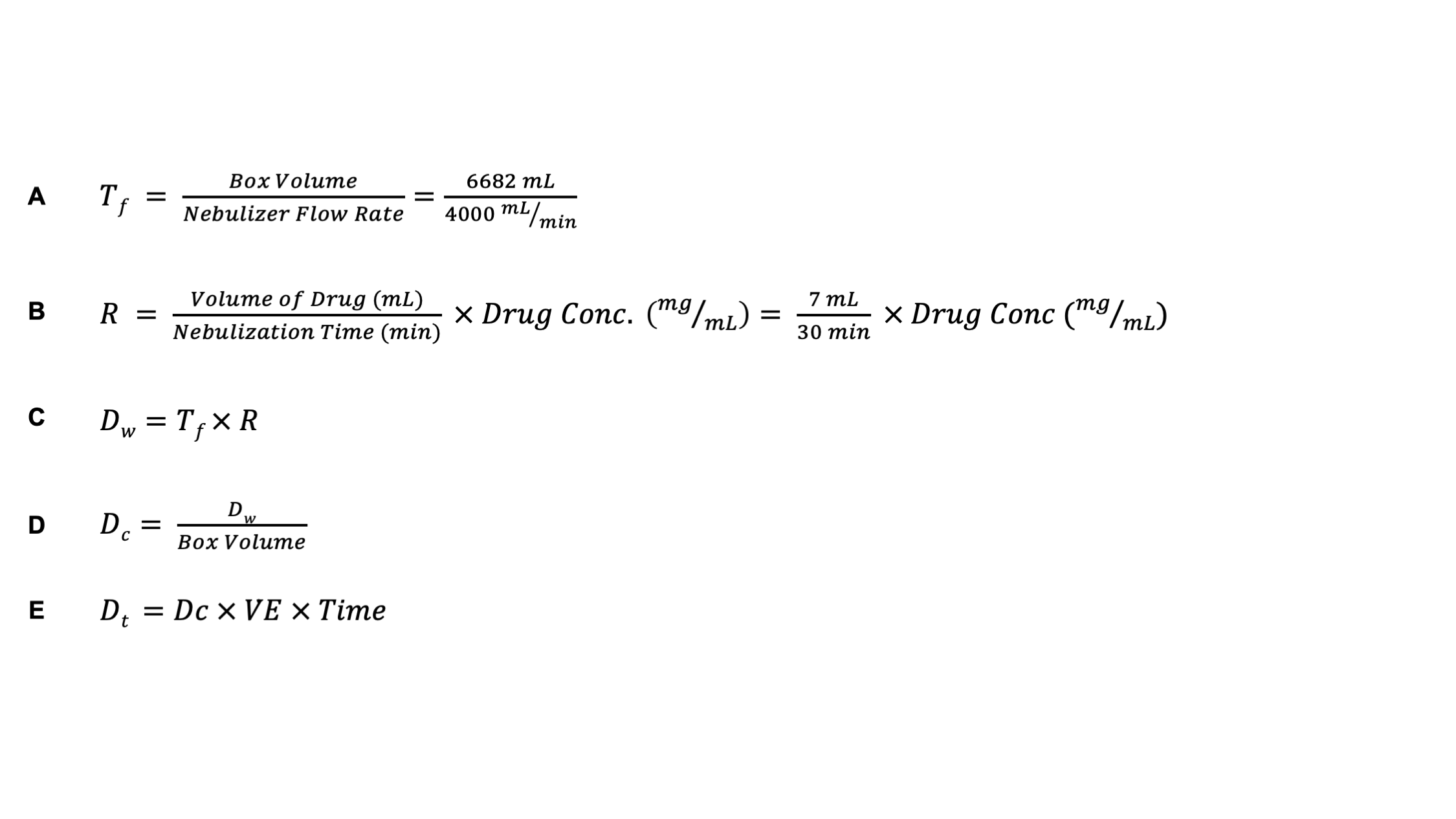
